# Topic: Hyper immune Bovine Colostrum as a Low-Cost, Large-Scale Source of Antibodies against COVID-19

**DOI:** 10.1101/2021.05.20.444932

**Authors:** Hassan Nili, Majid Bouzari, Hamid Reza Attaran, Nader Ghaleh Golab, Mohammad Rabani

## Abstract

Many different strategies have been used to fight against COVID-19 pandemic as a therapeutics or prophylaxis approaches. However, none of them so far have used, passive immune transfer using products from immunized farm animals. Hyper immune bovine colostrums (HBC) have been used against many different respiratory and gastrointestinal tracts infections during past decades.

Six mixed Holstein X Semental dairy cattle’s in their 6-7 months of gestation period years were chosen for hyper immunization with COVID-19 vaccine. An isolated and very well protected site was selected and equipped according to animal husbandry code of practice, used for animal experimentation. Specific IgG level against SARS-CoV-2 virus was measured before and after vaccination in the sera, and in the colostrum following parturition. Very high specific IgG level was detected one week following second vaccination in the sera and in first colostrums after parturition. Safety of the product was approved following phase 1 of clinical trials in 40 healthy volunteers. Phase 2 of the clinical trials is underway. Early results show effectiveness of the product in reducing sore throat and cough in early stages of SARS-CoV-2 infection.

## Introduction

The emergence of severe acute respiratory syndrome coronavirus 2 (SARS-CoV-2) has caused severe human respiratory infection (COVID-19) and has become more than global health crisis (1, 2). It had devastating effects on all aspects of life, from increasing family violence’s and abuses, to having catastrophic effects on world economy (3). On January, 2020, the World Health Organization announced that outbreak of SARS-CoV-2 is a public health emergency with international concern (4). Coronaviruses belong to the family of Coronaviridae, an enveloped virus with positive-stranded RNA (5). It contains four main structural proteins: spike (S) glycoprotein, envelope (E), membrane (M) and nucleocapsid (N). According to genetic analysis, SARS-CoV-2 is related to bats and pangolin coronavirus which place this virus in the Beta coronavirus, indicating that the origin of SARS-CoV-2 may be bats Coronavirus (BatCoV RaTG13), and pangolin may be the intermediate host (6, 7).

COVID-19 outbreaks showed that achievement of efficient vaccines are out of hand, especially in the early stages pandemic. Using animal models for production of large amount of specific antibodies could be used as an alternative approach against circulating pathogen during the pandemic, especially in immune compromised patients. Although passive immune transfer using convalescence plasma therapy have shown not to be clear in reducing mortalities in COVID-19 patients, there is no report on the efficacy of HBC in treatment of covid-19 patients(8). By hyper immunization of pregnant dairy cows in the late gestation periods using specific antigens, the concentration of specific immunoglobulins in the sera could be increase. Passive immunity from ingestion of colostrum and milk is essential for the survival of newborn animals. Therefore, it has been proved that vaccination of pregnant cows before calving can effectively prevent infection of newborn calves.

Antibodies level specially IgG1 will be reduced in the blood stream 2-3 wks before partition and actively will be transported through receptor mediated mechanism to the lacteal secretion, following parturition (9). The total amount of IgG1 obtained from each lactation could be as high as of 500 grams (9, 10). Oral HBC and milk not only can increase the mucosal immunity in the oral cavity, pharynx and upper respiratory tract of human, even could have immunomudulatory effects on the host immune system. Passive immunity caused by immunoglobulin transfer is a well known concept adopted by most mammals. IgG is one of the main components of immune activity found in milk and colostrums which can bind to many gastrointestinal and respiratory pathogens that infect humans such as Cryptosporidiosis, Shigellosis, Rotavirus, Respiratory Syncytial Virus (RSV), Human immunodeficiency Virus (HIV), Influenza, Enterotoxigenic *Escherichia coli*, and *Clostridium difficile* infection and support the cross-species activity of bovine and human IgG (11–19).

Immunoglobulins in breast milk are IgA, IgG1, IgG2 and IgM. On the contrary, IgG1 is the main immunoglobulin in cow’s milk, especially colostrum, while the concentration of IgM, IgA and IgG2 are lower (20). The concentration of IgG1 in colostrum is 100 times higher than milk (21). Beside specific antibodies, bovine colostrum contains many essential nutrients and bioactive components, including growth factor, immunoglobulin (Igs), lactoperoxidase (LP), lysozyme (Lys), lactoferrin (LF), cytokines, nucleosides, vitamins, peptides and oligosaccharides. These components are increasingly related to human health. IgG from unimmunized cattle can interact with different types of pathogens including viruses. Interestingly, bovine IgG can interact with the human Fcg receptor (FcgR), which can enhance antigen presentation to T-cells and phagocytosis of leukocytes (21, 22).

An experimental research showed that bovine colostrum increased the proportion of CD8+ T-cells after virus attack in mice(16). IgG also has other functions, including agglutinating pathogens, fixing complement to lysis pathogens, inhibiting pathogen metabolism by blocking enzymes, and neutralizing viruses. Bovine colostrum-derived IgG can inhibit the NF-kB signaling pathway and inhibit the production of pro-inflammatory cytokines in intestinal cells(23). Hyper immune bovine IgG can directly bind to the virus and prevent pathogen organisms from adhering to intestinal epithelial cells (17, 24, 25).

Consistent with this study, treatment of mice with immune IgG produced by cows immunized with LPS is associated with an increase in the number of NKT cells in the spleen, indicating that oral administration of hyper immune colostrum preparations can reduce chronic inflammation, including liver damage(26).

It has been reported that bovine IgG-derived colostrum is resistant to proteolysis, which supports the view that IgG-derived colostrum contains trypsin inhibitors, which can promote these antibodies survive throughout the gastrointestinal tract (27, 28).

In a group of high-risk cardiovascular patients, the effect of oral colostrum on hospital flu-related complications was studied. These patients received only colostrum, colostrum combination vaccination or single vaccination. Compared with the colostrum-only group, flu-related complications in the colostrum-only group were significantly reduced (29). In addition, colostrum prevents influenza infection in healthy volunteers at a rate equivalent to influenza vaccination (29).

Therefore, considering previous work on prophylactic and therapeutic effects of bovine colostrums derived immunoglobulin’s on different infectious organisms, we have used similar approach against COVID-19 infection. This research focuses on passive immunization and how hyper immune milk or colostrum collected from cows vaccinated with SARS-CoV-2 can be used to provide short-term protection against SARS-CoV-2 infection in humans. And can be used as an alternative method of immunization specially in more vulnerable individuals and as a prophylaxis in health care staffs. By this approach large scale and low cast production of immune components can be achieved to confront pandemic such as SARS-CoV-2.

## Results

### Clinical observation

Veterinary and laboratory check out of the pregnant animals and their fetus used in this experiment, did not show any abnormalities before and after vaccination. At least 3 doses of 2 ml vaccines were used for each animal immunization before parturition. No any changes were observed in the behavior and clinical signs such as body temperature, feed and water consumption of vaccinated pregnant animals. Also, adverse tissue reactions were not detected on the injection site in the thigh muscle after vaccination.

### ELISA results

As it is shown in table 1, three of the cattle had their forth vaccination before parturition. In three others, only three vaccine injection conducted, two of which delivered their calf few days after third vaccine injection. IgG level in the serum of six pregnant bovine raised following first vaccinations; however we observed highest IgG level increase one week after second vaccination (figure 1). Third and forth injection did not changed IgG level, compared to second vaccination.

**Figure 1:**
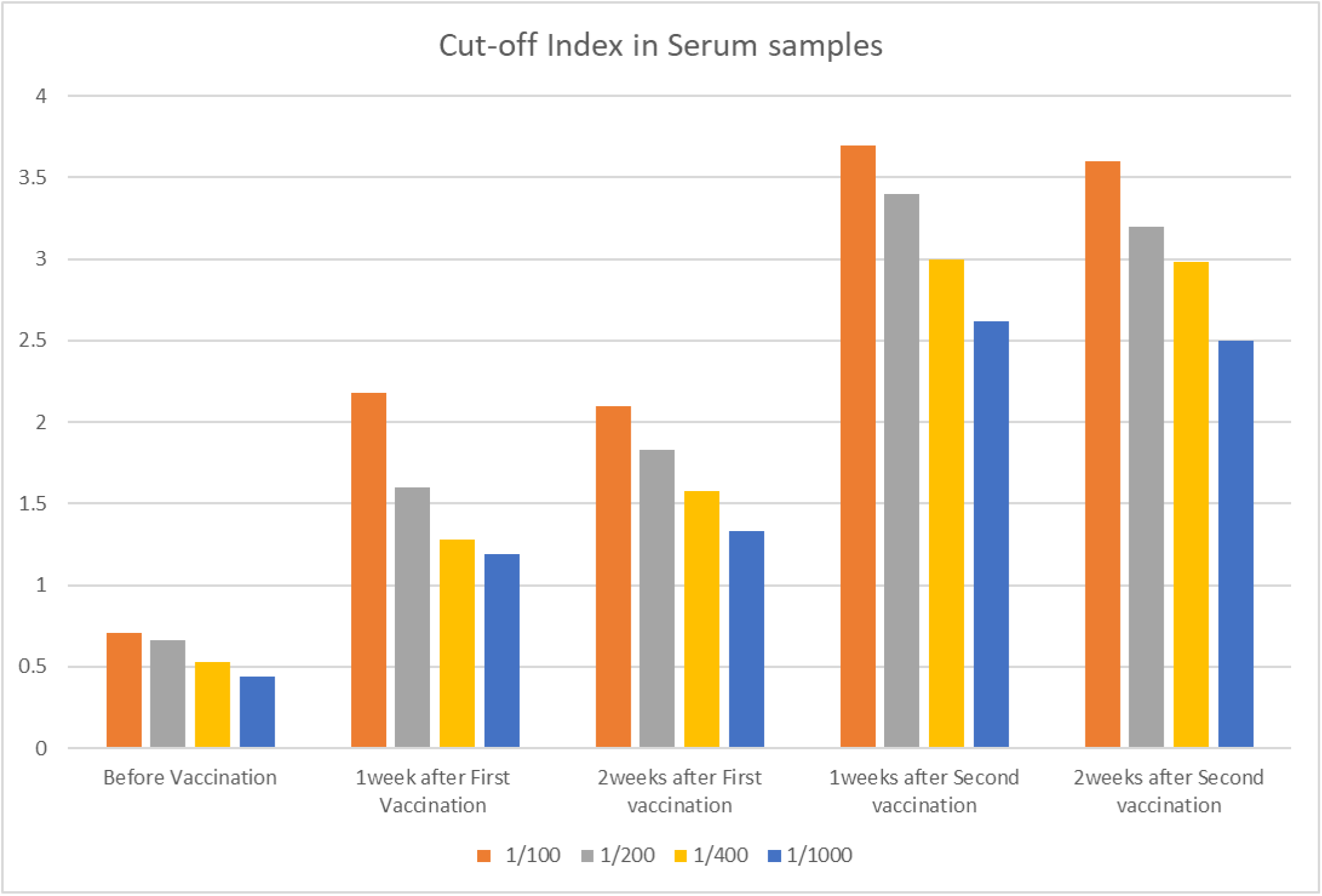
Mean specific IgG level in different serum dilutions of six pregnant cows before and after first and second vaccination, using ELISA test. Light absorbent by bovine IgG was measured at 450 & 630 wavelengths.

#### Virus neutralization

The virus was successfully neutralized by antibody present in the sera.

As it is shown in figure 2 high level of mean specific IgG is shown even in lowest dilution of the first colostrums (1/1000).

**Figure 2.**
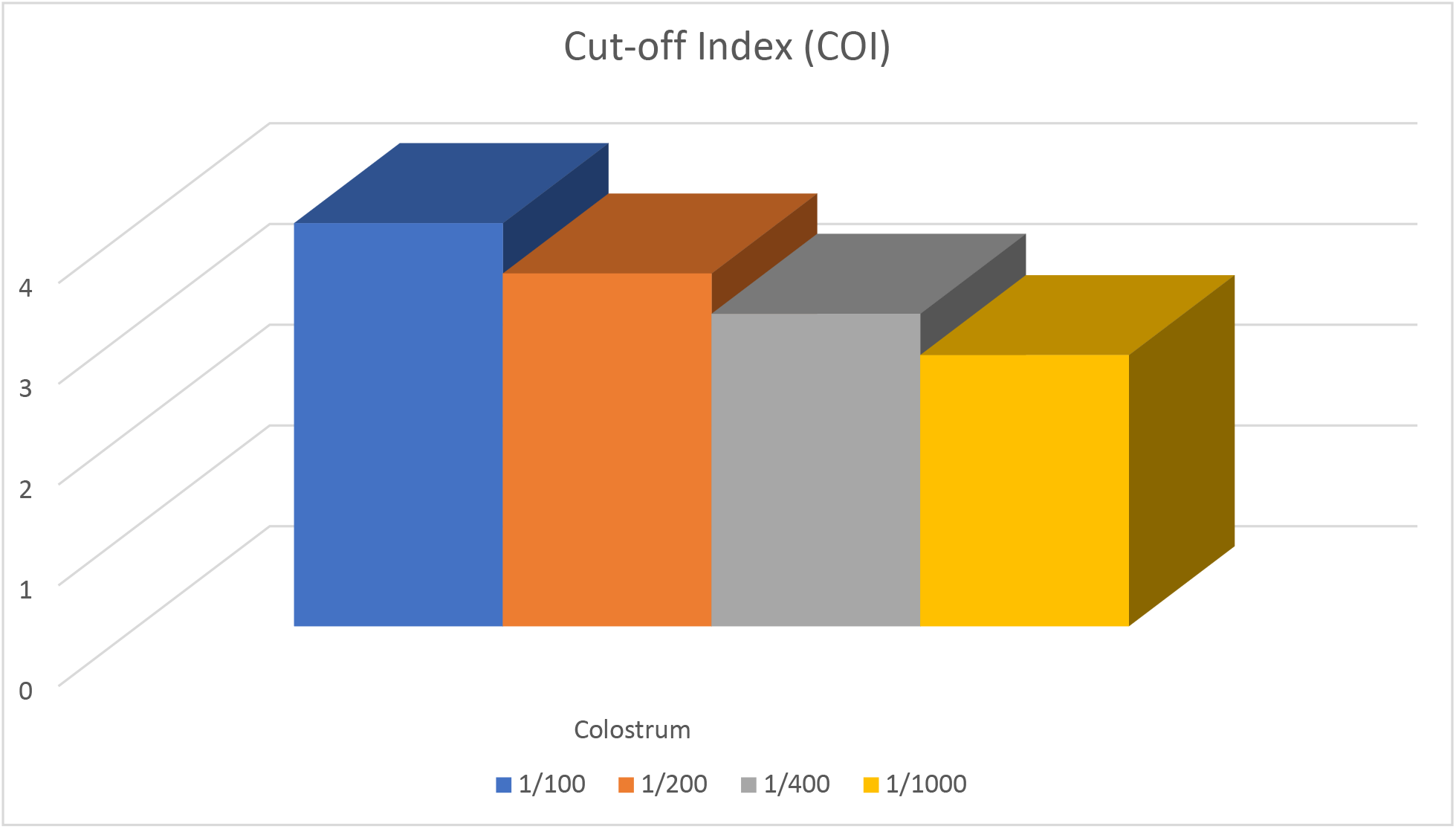
Mean specific IgG level in different colostrums dilutions of six cows, using ELISA test. Light absorbent by bovine IgG was measured at 450 & 630 wavelengths.

Comparison of dynamism of specific IgG level (mean) between serum collected before parturition, at parturition, first colostrums and milk obtained seven days following parturition in six pregnant cows are presented in figure 3. As indicated, there is a very sharp increase in specific IgG level against COVID-19 virus, following second vaccination. Just before parturition, IgG level sharply decreases in the serum and after parturition similar increase of IgG level has been observed in the first colostrum.

**Figure 3.**
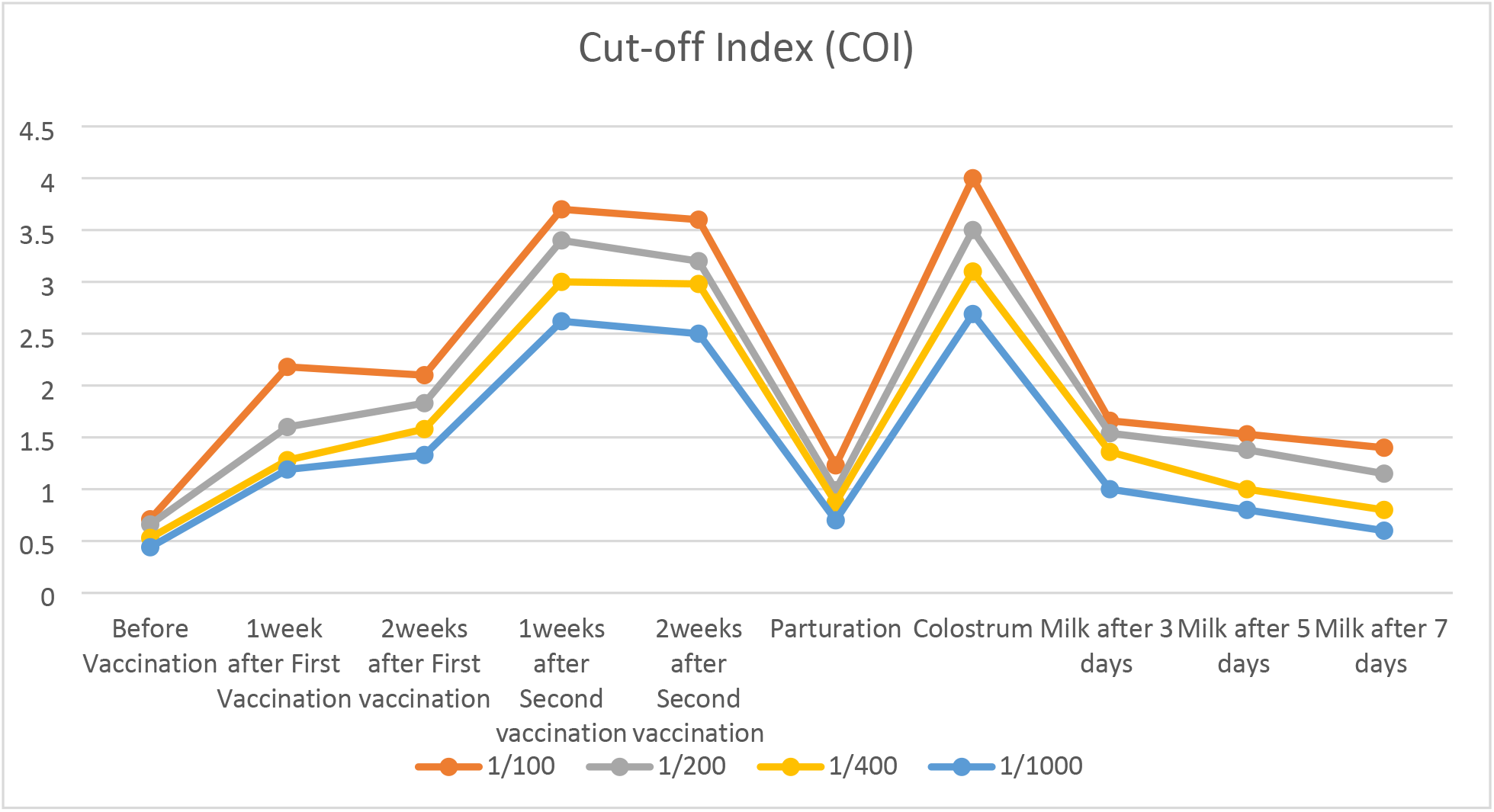
Dynamism of specific mean IgG level against covid-19 virus, in different dilutions, between serum collected before parturition, at parturition, first colostrums and milk obtained seven days following parturition in six pregnant cows, used in this study. Light absorbent by bovine IgG was measured at 450 & 630 wavelengths in ELISA test.

##### Clinical trial phase 1

there were no any adverse effects following frequent use of hyper immune bovine milk and colostrums in healthy volunteers, containing specific antibody against SARS covid-19 virus. Therefore the safety of the product was officially approved by ethic committee for research in biological sciences of Isfahan University of Medical Sciences.

##### Clinical trial phase 2

This clinical trial is still underway.

## DISCUSSION

Vaccinated pregnant cows behaved normally following vaccination and did not show any clinical signs and mortalities, there was no abortion, fever or changes in feed and water consumption, and any tissue reaction in the vaccination site. Although many FDA-approved and investigational anti-viral drugs, alone or in combination, are in use during the ongoing SARS covid-19 pandemic, none of the clinical trials so far have used bovine colostrums based immune components against COVID-19 (30). Considering the safety of the product which has been approved following conducting phase1 clinical trial during this study, immunomodulatory effects of the product can be studied in different stages of the disease and even as a prophylaxis, especially in health care workers, in front line of fighting covid-19 virus pandemic. Personal observations during last three months using hyper immune bovine milk and colostrums show that the products are efficacious especially in early stages of infection. Passive immune transfer using bovine immunoglobulin has been used in many respiratory and gastrointestinal tracts infections during last two decades. For more than 100 years it has been recognized that milk and maternal antibody provide passive immunity to a newborn infant via the transfer of bioactive factors and immunoglobulin’s (Igs) (31). Unique physiology of antibody transfer from mother to neonate in ruminants, which is through colostrum, has provided us, with massive amounts of antibody, immediately after parturition (9). This phenomenon provides very valuable immune components, with wide range of use in human health and medicine. By hyper immunization of pregnant cows in their late gestation period, we can increase the specificity of immune components that are available to us after parturition.

Perhaps because of this unique function, ruminant neonates are borne without Igs and 70-80 percent of total protein content in their colostrums is Igs (32, 33).

The results of current study show that the IgG level starts to decline 2-3 weeks before parturition, and this is because of active receptor mediated transfer of the antibody from blood stream to the mammary glands. These results are in agreement with previous research by Burton *et al* (9).

The level of IgG in the blood stream did not increased much after 3^rd^ and 4^th^ vaccination, this could be due to limited time period before parturition, as explained before, therefore in future study, we propose to use pregnant cow in their earlier gestation period, so there would enough time for evaluation of consequent antibody response 3^rd^ and 4^th^ vaccine injection.

High IgG level obtained even in the lowest dilution of the sera and colostrum, indicates that the vaccine used in this study has been able to show very good humeral immunity response in vaccinated cattles.

In addition to currently approved antiviral therapies, passive transfer of immune components through oral routs in dairy products could be an alternative strategy against the virus (16, 34). Although several new therapeutic strategies are emerging in these desperate times, none of them are based on specific bovine derived immunoglobulin’s.

Vaccination or passive immune transfer strategies both can be used in fighting COVID-19 infection. Each one has its own advantage and disadvantages,

Although vaccination is more popular and well-established strategy against infectious diseases, technical difficulties in conducting different clinical trials needed for achieving efficient, potent and safe vaccine are obstacles. Moreover, there is delayed antibody response after vaccination, considering the fact that the immune system of the host is functional and the response is appropriate. Using oral immune transfer strategy, will not have any contradiction with injecting vaccines, also the clinical trials for oral immune transfers are more practical and steadfast.

This idea dates back to the 1950s when Petersen and Campbell proposed that orally administered bovine colostrum from hyperimmunized cows could provide passive immune protection for humans(35).

Currently, the US Food and Drug Administration (FDA) have accepted the safety of hyperimmune milks on the basis of clinical studies that show no adverse health effects from these products(36, 37).

Phase one of the clinical trial was conducted to determine the safety of hyper immune bovine milk against COVID-19, using 150 ml on daily basis for up to 30 consequence days did not have any adverse effects in healthy volunteers aged between 18-65 years. In an ongoing phase II of clinical trial, some efficiency have been observed from anti-COVID-19 milk using 150ml of hyper immune milk.

**In conclusion,** In current COVID-19 and future pandemics, beside vaccination which is a very time consuming and complicated process, especially in the early stages, by passive immune transfer, using bovine hyper immune milk and colostrums large amount of specific abs could be available for prophylactic and therapeutic proposes (12, 13).

## Materials and Methods

Six mixed Holstein X Semental in their 6-7 months of gestation period aged between 3-4 years were chosen for hyper immunization with COVID-19 vaccine. Before purchase, cows were tested for Brucellusis and bovine Tuberculosis by local governmental veterinary organization. Also their health status and stage of gestation were examined by dairy farm veterinary specialist. An isolated and very well protected dairy farm was equipped and selected for the experiment in Zardanjan area in East of city of Isfahan-Iran. Animal were kept under close daily observation for adaptation to the new environment. An experienced animal husbandry engineer was employed for supervision of dairy cows during the experiment. Special Diet for dry period was purchased from Vahdat Company in the city of Isfahan.

### Vaccine preparation

This vaccine was produced according to the protocol of influenza vaccine production and FDA approved adjuvant was used as well. Briefly, the virus was isolated from the naso-pharynx samples taken from COVID-19 positive patients, cultured on **WHO** Vero cell line. The presence and purity of COVID-19 virus was checked by RT-PCR, Nano-Sensor and serum neutralizing tests(38). Each ml of the vaccine was contained 5 × 10 ^8.3^inactivated viral particles of COVID-19 virus. For viral inactivation formaldehyde was used according to the protocol. To check inactivating test, in proper laboratory animals’ model such as mouse and rat and Syrian hamster (39, 40) in groups of five, no virus was detected from pharyngeal swabs and blood samples for at least 2-month post inoculation, also no any clinical signs detected in all lab animals. Quality control tests for bacterial contamination such as blood agar, nutrient broth, thioglycollate broth, PPLO broth at 37°C, Sabouraud dextrose agar at 25°C were conducted on the harvested virus as well as vaccine.

### Hyper immunization

Animal experimentation was approved by ethic committee of the University of Isfahan. Animals were vaccinated with 2 ml of COVID-19 vaccine intramuscular in thigh muscle according to the vaccination schedule presented in the table 1.

**Table 1:** Vaccination time table of six pregnant mixed breed of dairy cattle in their late gestation period.

| Cow No | Vaccination date |  |  |  |
| --- | --- | --- | --- | --- |
|  | 1<sup>st</sup> | 2<sup>ed</sup> | 3<sup>rd</sup> | 4<sup>th</sup> |
| 1 | 14 Aug. | 28 Aug | 16 Sept | 25 Oct |
| 2 | 18 Aug | 1 Sep | 16 Sept | 25 Oct |
| 3 | 24 Sep | 10 Oct | 25 Oct | 25 oct |
| 4 | 11 Nov | 23 Nov | 9 Dec | N/A |
| 5 | 11 Nov | 23 Nov | 9 Dec | N/A |
| 6 | 11 Nov | 23 Nov | 9 Dec | N/A |

### Clinical observation

Before and after vaccination, pregnant cows were closely monitored on daily basis for any changes in behavior such as occasional systemic shock, itching, swelling or any adverse effects in the vaccination site in the thigh muscle. Also, they were monitored for any change in water and feed consumption, restlessness, increase in body temperature.

### Blood samples

All laboratory experiments were conducted in the Virology Research Center of University of Isfahan in conjunction with Zeitoon Isfahan Vaccine Innovators Company’s facilities. Blood samples were collected on weekly intervals from milk vein of the animal into special serum tubes, kept in room temperature for 30 min to coagulate red blood cells, and then centrifuged at 2500 g for 15 min at room temperature. Serum was removed and stored in aliquots at −20 C before use.

### Preparation of colostrum and purified colostrum IgG

HBC was collected immediately after parturition, quickly were pasteurized at 60°C for at least 60 mins. (Low temperature long time, LTLT). After pasteurization, the colostrums temperature brought to 4° C, then transferred to the laboratory, frozen at −20° C, until use.

### ELISA

Enzyme-linked immunosorbent assay (ELISA) was used for specific IgG measurement in the sera and supernatant obtained after centrifugation of colostrums after removal of fat and precipitant. Frozen colostrums were melt, centrifuged at 11000 g for 15-30 min at 4° C. protocols have been done according to ELISA Kit SARS-CoV-2 IgG (Pishtazteb, Iran). We just changed the secondary antibody and for this step horseradish peroxidase (HRP)-conjugated rabbit anti-bovine IgG antibody (Sigma) was used. In ELISA plate reader light absorbent by bovine IgG was measured at 450 & 630 wavelength. According to ELISA kit used in this experiment the Cut off value and Cut of index are obtained from the following formula.

> Cut off value = mean of light absorbent of negative controls + 0.15
>
> Cut off index (COI) = OD of sample/cut-off value

#### Viral neutralization test

Viral neutralization test was conducted in bio-safety level-3 laboratory at “Razi vaccine and Serum Research institute” of Iran.

##### Phase 1 of clinical trials

in at least 40 healthy volunteers, aged between 18-60 years, with no back ground medical complications has been conducted under the supervision of ethical committee for research projects in biological sciences with ethical code of IR.MUI.REC.1399.672, of Isfahan University of Medical Sciences. Each individual signed the consent form before entering phase 1 of clinical trials. Volunteers were given 150 ml of hyper immune bovine milk for up to 30 consequent days.

##### Phase 2 of clinical trials

A randomized double blind placebo controlled clinical trials including 40 hospitalized patients diagnosed by PCR and lung city scan to be positive for covid-19 in two hospitals affiliated to Isfahan University of medical Sciences is underway. Patients not having blood O_2_ saturation less than 90% were given 150 ml of hyper immune bovine milk or placebo twice on daily basis for 5 consequent days.

